# Affinity purification contaminants identified by cryo-EM and mass spectrometry

**DOI:** 10.64898/2026.03.19.712978

**Authors:** Emma R. Belcher, Steven W. Hardwick, Taiana Maia de Oliveira, Marko Hyvönen

**Affiliations:** Department of Biochemistry, University of Cambridge, 80 Tennis Court Road, Cambridge CB2 1GA, UK; Protein Sciences, Structure, and Biophysics, Discovery Sciences, R&D, AstraZeneca, Cambridge, UK

**Author notes:** Correspondence to: Marko Hyvönen.

**Keywords:** affinity purification, contaminants, Strep-tag, FLAG-tag, cryo-EM

## Abstract

Affinity chromatography is a powerful and therefore popular method for the purification of proteins for structural studies. The success of the technique relies on the specificity of the interaction between the target protein and the affinity resin. Here, we present the identification of two protein contaminants isolated from HEK293 cell lysate following affinity purification of twin Strep-tagged or FLAG-tagged proteins. The contaminants were identified as human propionyl-coenzyme A carboxylase (hPCC) and protein arginine methyltransferase 5 in complex with methylosome protein 50 (PRMT5:MEP50) via a combination of cryo-EM data processing and proteomic analyses. This report serves to illustrate how these contaminants may appear in cryo-EM datasets and to highlight the paramount importance of affinity chromatography resin specificity for efficient protein purification.

## Introduction

Single-particle analysis (SPA) by cryogenic electron microscopy (cryo-EM) has become a method of choice for determining the structures of proteins in near-atomic detail. Unlike X-ray crystallography, cryo-EM does not require protein crystallisation, making it particularly powerful for structural characterisation of proteins that are intrinsically unstable or difficult to produce in large quantities, such as membrane proteins. Because SPA relies on averaging many particles, the sample typically must be highly pure and homogenous in order to achieve high resolution reconstructions. Protein purification is therefore a critical step in the structure determination pipeline.

Conventional chromatography techniques separate proteins based on their physical or chemical properties, such as size, charge or hydrophobicity. While these classical methods can be used successfully for isolation of proteins from natural sources or from recombinant organisms after overexpression, their resolving power can be limited, as even the simplest of organisms contain thousands of proteins. Several sequential purification steps are therefore typically needed, resulting in lengthy workflows and reduced target protein yields.

Affinity chromatography is arguably the most powerful and widely used protein purification technique, enabled by recombinant DNA technologies to label target proteins with an affinity tag or fusion protein. The highly specific interaction between the target protein and its immobilised binder allows for efficient purification of the target protein from a complex mixture, such as cell lysate, with relatively high purity. This is especially effective for membrane proteins, which are typically expressed at low levels and are often unstable outside of their native membrane environment. The highly specific nature of affinity chromatography means that the target can often be isolated in a single purification step in a small volume.

Affinity tags and fusions come in many different varieties, from entire folded proteins like glutathione-S-transferase (GST) or maltose-binding protein (MBP), to short peptide tags that bind to a specific partner, such as the FLAG-tag, which binds to anti-FLAG monoclonal antibodies. The latter are more commonly used for purification as their small size means that fusion to the protein of interest is unlikely to affect its function. These epitope tags can be added easily to either the N-or C-terminus of the target protein sequence and used in combination for purification via orthogonal affinity chromatography steps or for downstream experiments. The use of these approaches relies on high affinity and specificity of the fusion partner and corresponding affinity matrix, to enable quantitative capture of the target proteins with minimal contamination. While that is generally the case, there are instances where the process is compromised by competing interactions of cellular components and this can have a significant impact on purity and yield of the target protein. Although noted in the literature, this is an issue with affinity purification that should be more widely appreciated.

We set out to isolate a multicomponent transmembrane receptor complex for structure determination by cryo-EM. The target was a tetrameric transforming growth factor (TGF)-β receptor signalling complex consisting of two copies of one type of receptor (ALK4) and two copies of another receptor (ACVR2A), together with their growth factor ligand, activin A. The receptor complex transduces the signal of growth factor binding across the plasma membrane. Full-length TGF-β receptor complexes have proved to be elusive targets for structure determination (1).

We produced the target complex in mammalian cells, with each type of receptor fused to a different affinity tag (1). The type I receptor ALK4 was fused to a twin Strep-tag for immobilisation on StrepTactin resins and detection by anti-Strep-tag antibodies in western blot analyses. The type II receptor ACVR2A was fused to a yellow fluorescent protein (YPet) to facilitate monitoring via fluorescence-detection size exclusion chromatography (FSEC), as well as a FLAG-tag for detection by anti-FLAG antibodies in western blot analyses. The protein component of the target complex (that is, excluding detergent) had an expected molecular weight of 316 kDa. Based on existing crystallographic data of the isolated extracellular domains (2) and AlphaFold3 models of the full-length receptors (1,3), we anticipated that the target complex would have a particle diameter of approximately 160 Å.

We purified the target complex by two different affinity chromatography approaches and applied the samples to cryo-EM grids. During data collection and processing, we detected populations of particles that did not match our expectations. Following careful data processing and proteomic analysis, we identified human protein contaminants of our affinity matrices that, despite being much less abundant in the sample than the target proteins, formed structurally homogeneous particles that dominated our cryo-EM datasets.

## Results

### StrepTactin resins enrich endogenously biotinylated proteins

In our first experimental approach, we immobilised the target complex via the twin Strep-tag on the type I receptor using StrepTactin affinity resin. StrepTactin is an engineered form of streptavidin that binds tightly to the Strep-tag peptide, enabling specific capture of a target protein fused to this tag (4). A key benefit of this affinity chromatography approach is that bound proteins can be eluted from the StrepTactin resin under gentle physiological conditions by competition with *d*-desthiobiotin.

In our experiments, the type I receptor was captured by the resin, and the type II receptor was also detected in the elution fraction by western blot analyses, suggesting successful purification of the entire receptor complex. The elution was purified further by fluorescence size exclusion chromatography (FSEC). Fractions of the expected retention volume containing fluorescent signal from the type II receptor component were frozen on cryo-EM grids.

During data collection and ‘on the fly’ preprocessing, a 3D volume with D3 symmetry was reconstructed from just 3097 particles and refined to 6.23 Å **(Figure 1)**. This symmetry was unexpected for the target complex, which was anticipated to be tetrameric. In addition, the target complex was expected to be compositionally and conformationally heterogeneous. The reasonably high resolution of the volume generated from a low number of particles suggested that this represented a rigid, homogenous protein or complex, not matching our expectations for the target protein complex. We searched the EMPIAR EMDB for entries matching this symmetry and provisionally identified the volume as human propionyl-coenzyme A (CoA) carboxylase (hPCC; EMD-33729; PDB:7ybu (5)).

**Figure 1.**
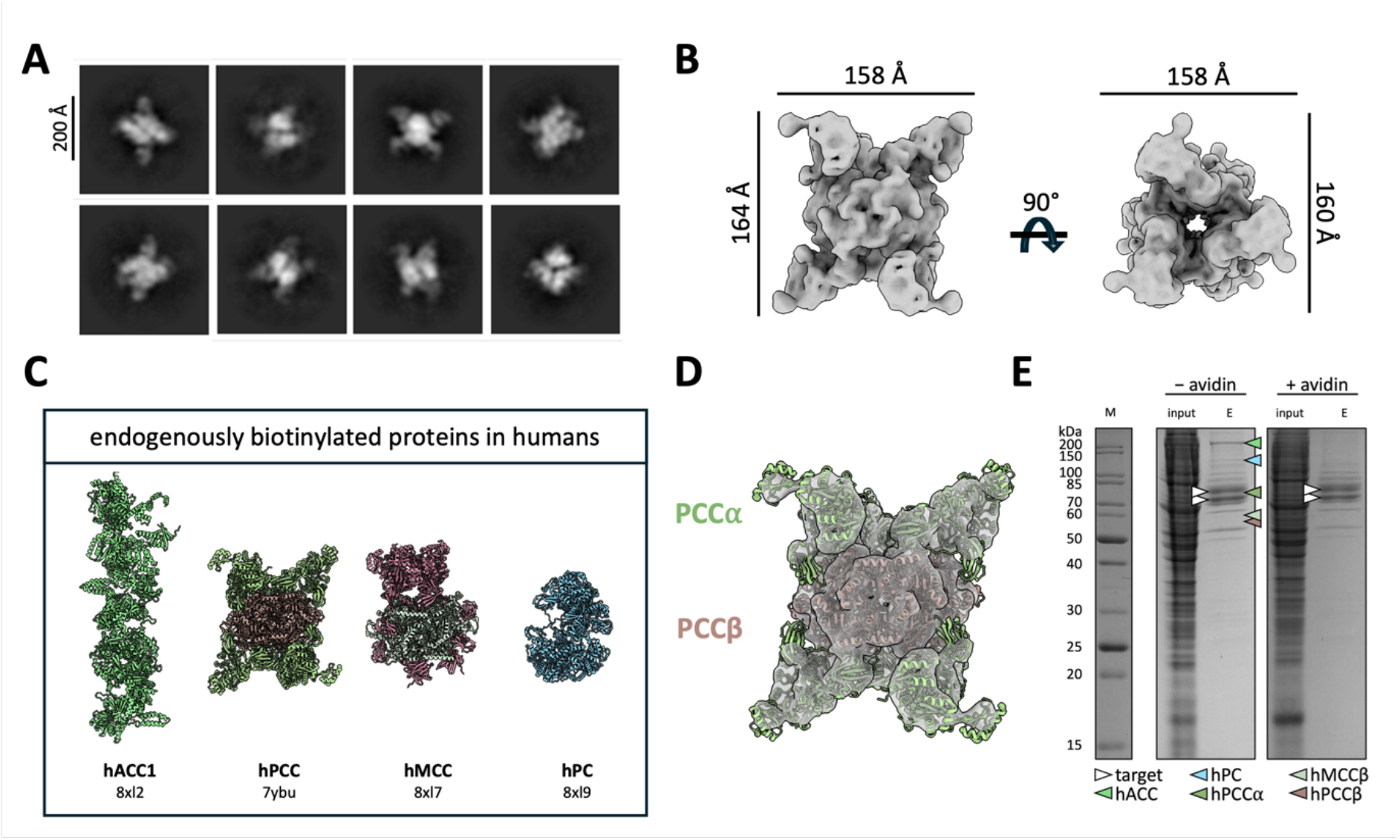
Human propionyl-coenzyme A (CoA) carboxylase (hPCC) was identified as a contaminant of StrepTactin resin by cryo-EM. (A) Selected 2D classes illustrate how the contaminant appeared during cryo-EM data preprocessing. (B) A 3D volume with a reported resolution of 6.23 Å was generated from 3097 particles. The approximate dimensions of the volume (measured along its principal axes) are shown. (C) Proteomic analysis identified all four endogenously biotinylated human proteins as hits in the sample. Cartoon representations of human acetyl-CoA carboxylase (hACC1; PDB:8xl2), hPCC (PDB:7ybu), human methylcrotonyl-CoA carboxylase (hMCC; PDB:8xl7) and human pyruvate carboxylase (hPC; PDB: 8xl9) are shown. (D) An atomic model of hPCC (PDB:7ybu), consisting of six chains of PCCα and six chains of PCCβ, fits well into the generated volume. (E) A Coomassie-stained gel shows the elution fractions after purification of a preparation of the type II receptor construct (85 kDa doublet band), with faint bands corresponding to biotinylated contaminants identified during proteomic analysis indicated with arrowheads. Addition of 0.5 mg/mL avidin to the sample during solubilisation successfully eliminated biotinylated contaminants. M: marker, E: elution.

hPCC is a biotin-dependent mitochondrial enzyme that plays a role in metabolism (5,6). The holoenzyme consists of six chains of each of two subunits, PCCα and PCCβ. The α-subunit contains a biotin carboxylase domain and a biotin carboxyl carrier protein domain, meaning that hPCC is biotinylated during its catalytic cycle. hPCC is one of four biotinylated proteins found in human cells — acetyl-CoA carboxylase, pyruvate carboxylase and methylcrotonyl-CoA carboxylase complete the set (7,8). This biotinylation explains the presence of hPCC in our sample, as biotin interacts almost irreversibly with StrepTactin. In fact, existing structural models of these endogenously biotinylated proteins were obtained by purifying untagged proteins on the StrepTactin resin used here (9,10). These contaminants were therefore being enriched by the StrepTactin affinity chromatography resin, competing with the Strep-tagged receptor complex for binding. Proteomic analysis revealed that, in addition to the target receptors, all four endogenously biotinylated human proteins were present in the sample, with the volume generated during cryo-EM data processing clearly matching existing structural models of hPCC by visual inspection **(Figure 1D)**. While the other endogenously biotinylated proteins were present in the sample, the diameter used for particle picking and box size for extraction, which were chosen based on the expected dimensions of the target complex, may have resulted in particles of hPCC being picked preferentially during data processing.

While the contamination of StrepTactin resins with biotin during purification from mammalian cells is expected, the extent and repercussions of co-purification of biotinylated protein contaminants are not widely appreciated and, for example, product manuals for these affinity matrices do not raise this as an issue. Fortunately, there are solutions to overcome this problem. Biotinylated proteins can be blocked from binding to StrepTactin by the addition of avidin, which binds extremely tightly to biotin, but does not bind the Strep-tag (11). Biotinylation sites are therefore ‘masked’ by the avidin, and biotinylated proteins can no longer interact with StrepTactin. In our experiments, addition of avidin to the sample prior to incubation with the resin successfully eliminated biotinylated contaminants **(Figure 1E)**. However, it also caused precipitation in samples, leading to insufficient yield of our target protein for downstream applications. Adding an excess of recombinant avidin to the preparation also added to the cost of the purification significantly. We therefore switched to a different purification strategy.

### Protein arginine methyltransferase 5 is non-specifically enriched by anti-FLAG M2 resin

In our target complex, the type II receptor was expressed with a FLAG-tag, a highly charged 8-residue peptide epitope, for detection by western blot. Conveniently, this tag also enabled us to purify the target complex using a different affinity chromatography approach, using an anti-FLAG monoclonal antibody-conjugated resin. In this experimental approach, the type II receptor is immobilised on anti-FLAG M2 resin, with the type I receptor co-eluting from the resin as part of the complex. Again, bound proteins can be eluted under gentle physiological conditions, by competition with a 3XFLAG peptide.

Receptor complex purification was confirmed by co-elution of the Strep-tagged type I receptor with the type II receptor during purification on anti-FLAG M2 resin. After size exclusion chromatography, the sample was analysed by cryo-EM. Similarly to the sample from the Strep-tag purification, several 2D classes containing relatively few particles (around 10000 in total) emerged with clear features. These classes were used to reconstruct a 3D volume with a resolution of 4.53 Å **(Figure 2)**. We suspected that this was another contaminant, due to the small number of particles required to reach this resolution. Mass spectrometric analysis confirmed that this contaminant was protein arginine methyltransferase 5 in complex with methylosome protein 50 (PRMT5:MEP50). Again, we could match our 3D volume to the published structure of this complex (EMD-27078; PDB:8cyi (12)).

**Figure 2.**
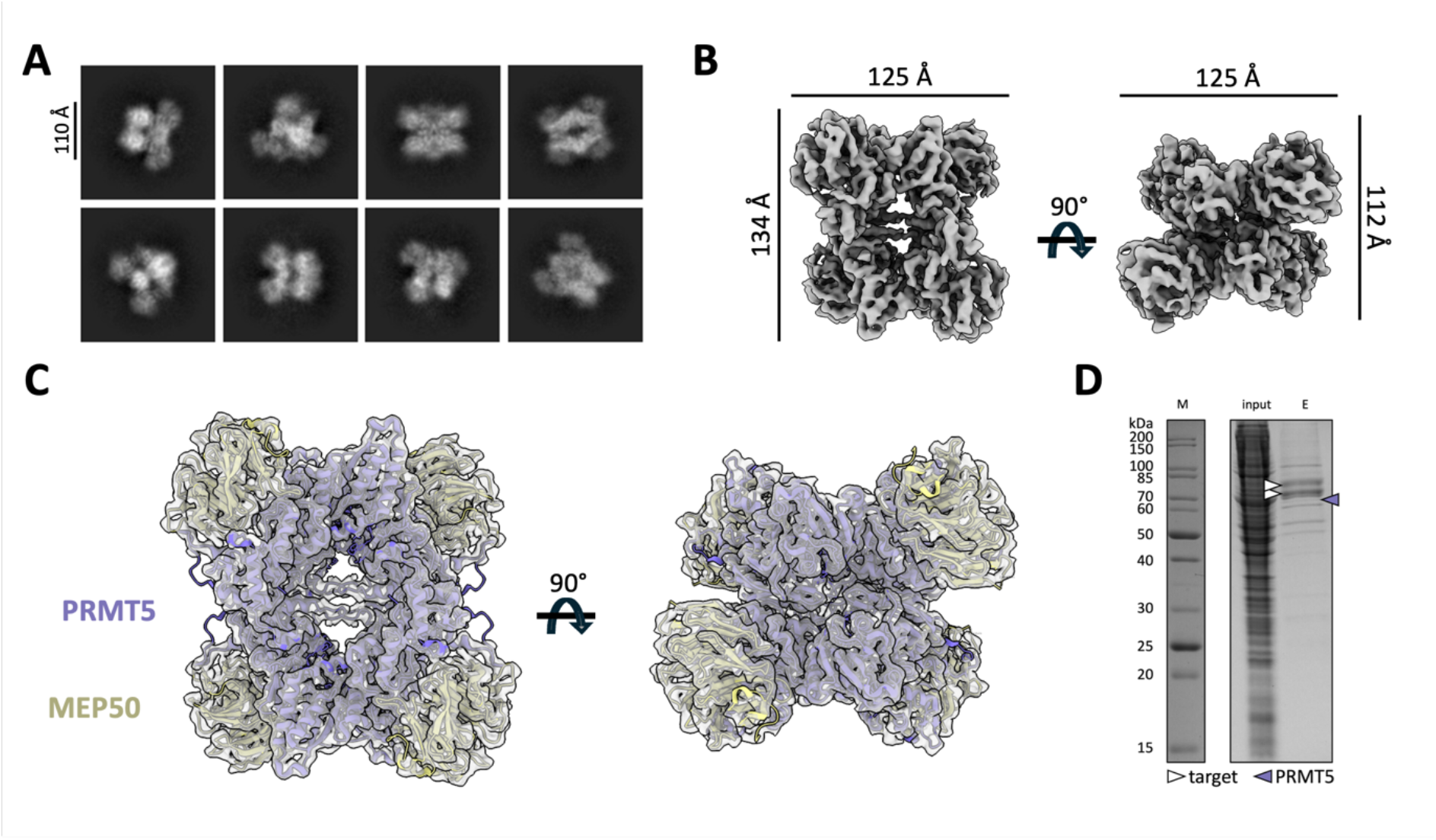
Protein arginine methyltransferase 5 in complex with methylosome protein 50 (PRMT5:MEP50) was identified as a contaminant of anti-FLAG M2 resin by cryo-EM. (A) Selected 2D classes illustrate how the contaminant appeared during cryo-EM data preprocessing. (B) A 3D volume with a reported resolution of 4.53 Å was generated from 9243 particles. The approximate dimensions of the volume (measured along its principal axes) are shown. (C) Four copies of an atomic model of PRMT5:MEP50 (PDB:8cyi) fit well into the generated volume. (D) A Coomassie-stained gel shows the elution fractions after purification of a preparation of the type II receptor construct (85 kDa doublet band), with a faint band corresponding to PRMT5 (expected molecular weight: 73 kDa) indicated with a purple arrowhead. MEP50 (33 kDa) was also identified by LC-MS/MS, but is not visible on the gel, likely due to low concentration. M: marker, E: elution.

PRMT5:MEP50 is a protein complex that plays a regulatory role in transcription and is therefore involved in many important cellular processes (13). In humans, the PRMT5:MEP50 complex consists of four copies of the methyltransferase PRMT5 and four copies of MEP50, which is thought to act as an adaptor protein for a range of protein substrates. Although PRMT5 contamination in anti-FLAG purifications has been noted elsewhere (14–16), it is not clear how PRMT5 is enriched by the anti-FLAG M2 antibody as it does not contain the FLAG-tag sequence. The surface of the complex is relatively charged (13), so it may be that part of the surface mimics the highly charged FLAG peptide sequence. The structure of the anti-FLAG M2 Fab fragment bound to the FLAG peptide provides a molecular basis for the recognition of the peptide tag by the anti-FLAG M2 antibody via multiple salt bridges and hydrogen bonds (PDB:8rmo (17)). We used AlphaFold3 to model the Fab fragment of the anti-FLAG M2 antibody bound to the PRMT5:MEP50 complex **(Figure 3)** (3). While the PRMT5:MEP50 complex and the Fab complex were each predicted with high confidence, the interaction interface between the Fab and PRMT5 was not reliably predicted by AlphaFold3 (mean predicted aligned error (PAE) of close to 30 Å). Nonetheless, the five models generated by AlphaFold3 show anti-FLAG M2 Fab bound to PRMT5 at a non-linear surface epitope, which would be consistent with the fact that we have not detected PRMT5 in western blots with anti-FLAG antibodies, where non-linear epitopes would be lost as a result of protein denaturation during SDS-PAGE. This speculative interaction interface has yet to be validated experimentally.

**Figure 3.**
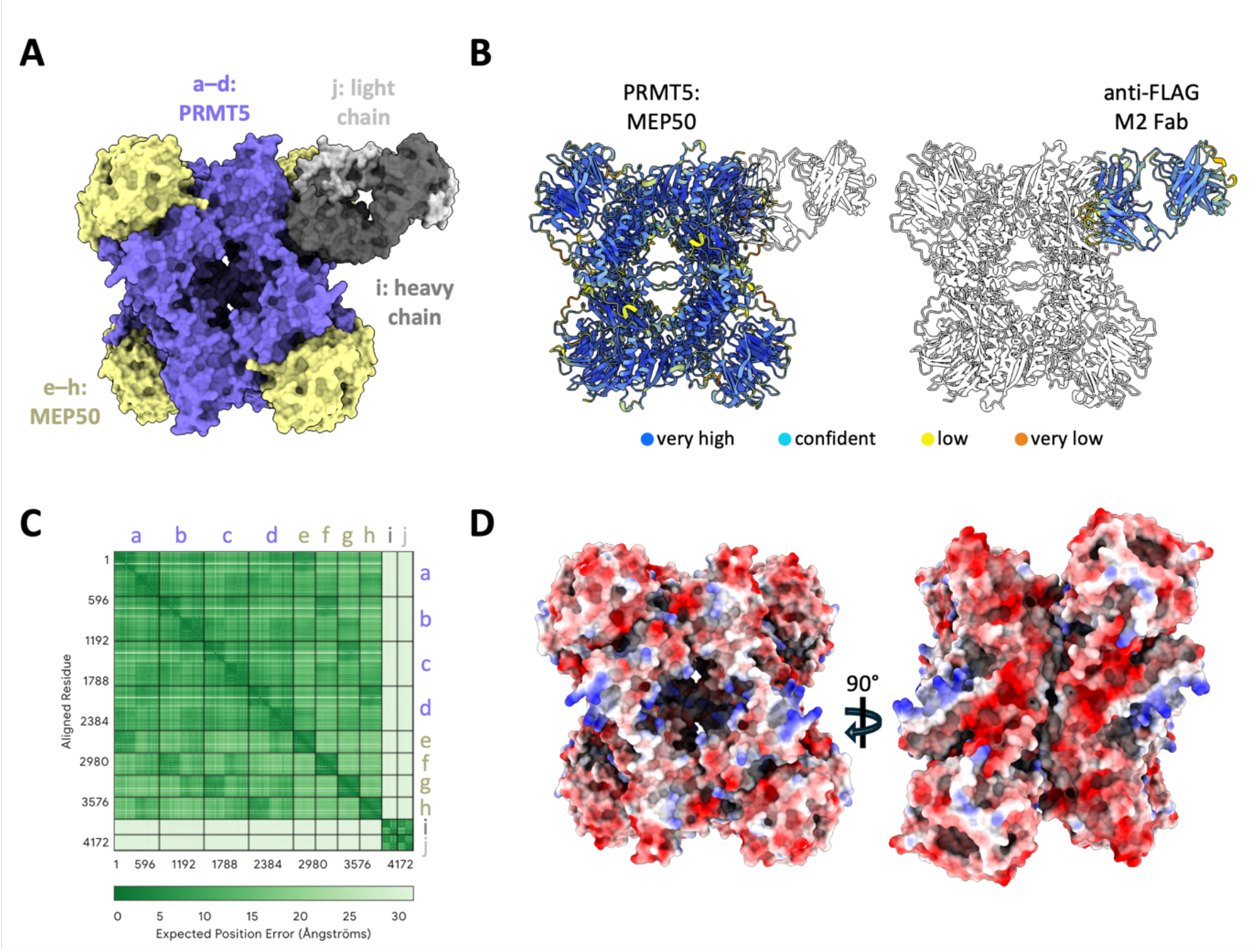
AlphaFold3 prediction of anti-FLAG M2 Fab bound to PRMT5:MEP50 suggests that the Fab may interact with PRMT5 at a non-linear surface epitope. (A) AlphaFold3 was used to generate a model of 4 chains of PRMT5 (purple), 4 chains of MEP50 (yellow) and the anti-FLAG M2 Fab heavy (dim grey) and light (light grey) chains (17). The overall predicted structure had an ipTM score of 0.78 and a pTM score of 0.81. (B) The PRMT5:MEP50 complex and the Fab complex were each predicted with very high confidence, with pLDDT > 90 (dark blue) for most residues. (B) The chains of the PRMT5:MEP50 complex were positioned with reasonable confidence relative to each other (mean predicted aligned error (PAE) of 12 Å). Similarly, the relative position of the Fab chains was reasonably confidently predicted (mean PAE of 8 Å). However, the interaction between the Fab (chains i and j) and PRMT5 (chain a) is not confidently predicted, with high mean PAE of around 30 Å. (C) The surface of PRMT5:MEP50 is highly charged, which may explain its recognition by the antibody against the highly charged FLAG-tag.

## Discussion

We have presented here how efforts to isolate a doubly affinity-tagged protein complex from mammalian cells resulted in enrichment of protein contaminants by two distinct affinity chromatography resins. These contaminants were first detected during cryo-EM data processing, and their identities were verified by mass spectrometry. Their enrichment reduced the effective binding capacity of the resins for the target complex, with endogenous proteins competing directly with the target proteins for resin binding. Furthermore, the relative structural homogeneity of the contaminants, compared with the flexibility of the target complex, exacerbated these challenges during cryo-EM data processing, as non-target particles dominated 2D class averages.

Affinity resins are expensive reagents and can represent a limiting factor in many projects, even in the absence of the additional complications described here. We report these findings to highlight how popular, commercially available affinity resins – often perceived as highly specific – can nonetheless enrich contaminant proteins. Contamination during affinity purification is not a new observation. Arguably the most popular affinity chromatography approach relies on the interaction between His-tags and an immobilised metal-ion chromatography (IMAC) resin like Ni-NTA. This purification approach is well known for its limited specificity and for enriching host proteins when suboptimal conditions are used. Common IMAC contaminants have been extensively catalogued (18), and numerous strategies have been developed to mitigate these effects when purifying recombinant proteins from bacterial systems. These include engineered *E. coli* strains in which common contaminant proteins are modified to reduce their affinity for chromatography resins or to enable their removal via orthogonal affinity purification steps (19). Such approaches are unlikely to be feasible in mammalian expression systems. However, alternative strategies – such as the inclusion of avidin during Strep-tag purification – can help mitigate contamination.

Beyond highlighting issues of resin-derived contamination, our findings also illustrate how such contaminants manifest in cryo-EM datasets and can disproportionately influence downstream analysis. With recent advances in cryo-EM throughput and achievable resolution, these issues have become increasingly apparent. While similar contaminants may have been present in samples prepared for crystallographic studies, they were likely at insufficient concentrations to crystallise independently. In contrast, even when present as minor components in cryo-EM samples, homogeneous contaminants can be readily detected during image processing. In our case, the intrinsic flexibility of the target complex likely allowed the comparably rigid contaminants to dominate in single-particle analysis, despite their relatively low abundance in the final sample.

We were able to eliminate the identified contaminants by combining the two affinity purification steps, albeit at the cost of reduced target protein recovery due to competition during the first purification step. Looking forward, the development of highly specific affinity chromatography reagents, such as nanobody-based affinity tools, or engineered resins that do not react with cellular components, such as HaloLink resins, may provide improved specificity. Such approaches could enable efficient immobilisation of target proteins while maintaining both high yield and high purity.

This note serves as a reminder that affinity chromatography resins, while powerful, are not perfectly selective and that contaminant enrichment can compromise both purification efficiency and cryo-EM data analyses. As cryo-EM continues to push toward higher resolution and throughput, particularly as the first-choice method for structure determination of protein targets that are considered ‘difficult’ due to low expression levels, instability or flexibility, improved affinity purification strategies will be essential to avoid trade-offs between target yield, purity, and data quality.

## Materials and methods

For detailed methods, see *PhD thesis, E. R. Belcher* (1).

### Protein expression and solubilisation

Human embryonic kidney Freestyle 293-F (HEK293F) cells were maintained at 37 °C with 70% humidity and 8% CO_2_ in air, with shaking at 125 rpm. Cells were passaged twice weekly. Cells were transfected with constructs encoding ALK4-ST (twin Strep-tag sequence: SAWSHPQFEKGGGSGGGSGGSAWSHPQFEK) and ACVR2A-YPet-FLAG (FLAG-tag sequence: DYKDDDDK). For a 500 mL culture, a total of 750 μg plasmid DNA and 1.125 mL 1 mg/mL polyethylenimine (PEI, Polysciences, 23966), pH 7, were added to 50 mL cells at 1 × 10^7^ cells/mL in a 2 L flask (Corning, 431255). After incubation for 1 hour, cells were diluted with Freestyle media to a total volume of 500 mL at 1 × 10^6^ cells/mL. Target proteins were expressed for 3 days. Cells were harvested by centrifugation and cell pellets were used immediately for purification or stored at −20 °C.

Cells from a 500 mL culture were lysed in 25 mL solubilisation buffer (100 mM Tris-HCl, pH 8, 150 mM NaCl, cOmplete protease inhibitors (Merck, 11836170001), 1% (m/v) dodecyl-β-D-maltoside supplemented with 0.1% (m/v) cholesterol hemisuccinate (DDM/CHS; Thermo Scientific, A50942)) for 2 hours at 4 °C with gentle mixing by rotation. After centrifugation at 21000 *g* for 15 minutes, clarified lysate was subjected to affinity purification.

### Purification on StrepTactin resin

For cryo-EM data presented here **(Figure 1)**, the sample was purified using StrepTactin-Sepharose resin (IBA Lifesciences, 12846436). Clarified lysate was incubated with resin, pre-equilibrated in wash buffer (100 mM Tris-HCl, pH 8, 150 mM NaCl, 0.02% DDM /0.002% CHS), for 1 hour at 4 °C with gentle mixing by rotation. Proteins were purified by gravity flow as per the manufacturer’s instructions. Proteins were eluted in elution buffer (10 mM *d*-desthiobiotin, 100 mM Tris-HCl, pH 8, 150 mM NaCl, 0.02% DDM /0.002% CHS) and concentrated using a spin concentrator with a 100 kDa molecular weight cut-off. Similar 3D volumes were obtained from preparations using StrepTactin-MagBeads (Cube Biotech, 34225).

### Purification on anti-FLAG M2 resin

Clarified lysate was incubated with anti-FLAG M2 resin (Merck, A2220), pre-equilibrated in wash buffer (100 mM Tris-HCl, pH 8, 150 mM NaCl, 0.02% DDM /0.002% CHS), for 1 hour at 4 °C with gentle mixing by rotation. Proteins were purified by gravity flow as per the manufacturer’s instructions. Proteins were eluted in elution buffer (0.1 mg/mL 3XFLAG peptide (GenScript, RP21087), 100 mM Tris-HCl, pH 8, 150 mM NaCl, 0.02% DDM /0.002% CHS) and concentrated using a spin concentrator with a 100 kDa molecular weight cut-off.

### Purification by fluorescence-detection size exclusion chromatography

After affinity purification, pooled elution fractions were further purified by size exclusion chromatography on a Superdex 200 Increase 10/300 GL column (Cytiva, 28990944) on an ÄKTA pure system (Cytiva) connected to a RF-20A fluorescence detector (Shimadzu). YPet (fused to type II receptor ACVR2A) was excited at 517 nm and emission was measured at 530 nm. Fractions of the expected retention volume containing fluorescent signal were pooled, concentrated using a spin concentrator with a 100 kDa molecular weight cut-off, and used immediately for cryo-EM.

### Cryo-EM grid preparation

Grid preparation was carried out in the BiocEM Cryo-EM Facility (Department of Biochemistry, University of Cambridge). The human propionyl-coenzyme A (CoA) carboxylase (hPCC) dataset was collected on a Holey carbon grid supported by gold mesh (Quantifoil, Au R1.2/1.3 300 mesh). The dataset containing protein arginine methyltransferase 5 in complex with methylosome protein 50 (PRMT5:MEP50) was collected on a Holey carbon grid supported by copper mesh (Quantifoil, Cu R1.2/1.3 200 mesh). Grids were glow discharged using a PELCO easiGlow system (Ted Pella, Inc) for 1 minute at a current of 25 mA. Using a Vitrobot (Thermo Fisher Scientific) set to 4 °C and 95% humidity, 3–4 μL sample were applied to the grid. After blotting for 3 seconds, the grid was plunge-frozen in liquid ethane. Grids were stored in liquid nitrogen and clipped before screening.

### Cryo-EM data collection and processing

#### Dataset containing hPCC

The dataset containing hPCC was collected on a Thermo Scientific Talos Arctica with a Falcon 3EC detector operating with an acceleration voltage of 200 kV (BiocEM Cryo-EM Facility, Department of Biochemistry, University of Cambridge). Data were collected at a nominal magnification of 73000X, pixel size of 1.37 Å/px, and a total dose of 58.88 e^−^/Å^2^.

During data collection, ‘on-the-fly’ data preprocessing was conducted using Warp (20). Particles were extracted with a box size of 300 pixels. A 3D volume, generated from 3097 particles, refined to 6.23 Å with D3 symmetry imposed. An atomic model of hPCC (PDB:7ybu (10)) was fitted into this volume using the ‘Fit in Map’ function in ChimeraX-1.9 (21).

### Dataset containing PRMT5:MEP50

The dataset containing PRMT5:MEP50 was collected on a Thermo Fisher Scientific Titan Krios with a bottom-mounted Falcon 4i detector operating with an acceleration voltage of 300 kV (Cambridge Pharmaceutical Cryo-EM Consortium, within the Department of Materials Science & Metallurgy, University of Cambridge). Movies were collected in counting mode in compressed TIF format at a nominal magnification of 120000X, pixel size of 0.654 Å/px, and a total dose of 60 e^−^/Å^2^.

The dataset containing PRMT5:MEP50 was processed in CryoSPARC (22) **(Figure S1)**. After patch motion correction and CTF estimation, 1060287 particles of diameter 200–400 were picked from 12052 micrographs using the Blob Picker tool. Particles were extracted from micrographs using a box size of 400 pixels and subjected to 2 rounds of 2D classification, resulting in a set of 10778 particles. Three 3D volumes were generated *ab initio* and the largest class, containing 6998 particles, underwent non-uniform refinement. This volume had a reported resolution of 4.88 Å. In order to search for other particles of this suspected contaminant, the particles and micrographs contributing to this volume were used to train a Topaz model for particle picking (23), using an estimated particle diameter of 130 Å. The Topaz model was used to pick 18476 particles which were extracted with a box size of 400 pixels. After 2D classification into 50 classes using 40 online-EM iterations and a batch size of 1000 per class, 3 volumes were generated *ab initio* with a maximum resolution of 6 Å and initial resolution of 25 Å. After non-uniform and local refinement of the largest 3D class, 9896 particles contributing to the resulting volume were classified into 15 2D classes, with a maximum resolution of 4 Å and ‘force max over poses/shifts’ set to false. The 9243 particles contained within 12 selected 2D classes were used as the input for non-uniform and local refinement of the volume. Gold-standard Fourier shell correlation (GSFSC) curves with a threshold of 0.143 indicated that the final volume, generated from 9243 particles, had an overall resolution of 4.53 Å. An atomic model of PRMT5:MEP50 (PDB:8cyi (12)) was fitted into this volume using the ‘Fit in Map’ function in ChimeraX-1.9 (21). Three additional copies of this model were generated and fitted into the volume individually to illustrate the D2 point symmetry of the biological assembly. Refining the volume with D2 symmetry imposed improved the reported resolution to 4.11 Å.

### AlphaFold3 structure prediction and analysis

The structure of the anti-FLAG M2 Fab bound to the PRMT5:MEP50 complex was predicted using AlphaFold3 (3,17). The PRMT5:MEP50 complex was first predicted alone, using the full-length protein sequences retrieved from UniProt (PRMT5: O14744; MEP50: Q9BQA1). The N-terminal region of PRMT5 and the N- and C-terminal regions of MEP50 were predicted to be highly flexible. These regions were removed for the prediction including the Fab, in line with the sequences provided in the structural model of PRMT5:MEP50 (PDB:8cyi). The structure of the anti-FLAG M2 Fab bound to the PRMT5:MEP50 complex was predicted using four copies of PRMT5 (residues 13–637), four copies of MEP50 (residues 21–329), one copy of the anti-FLAG M2 Fab heavy chain (residues 1– 217 per PDB: 8mro), one copy of the anti-FLAG M2 Fab light chain (residues 1–219 per PDB:8rmo) and one chloride ion. The five models generated by AlphaFold3 were visualised and analysed in ChimeraX-1.9 (21). The mean predicted aligned error (PAE) was estimated in PAE viewer (24).

### Proteomic analysis

The presence of the target proteins and identity of suspected contaminants was confirmed by liquid chromatography with tandem mass spectrometry (LC-MS/MS). Proteomic analysis was conducted by the Cambridge Centre for Proteomics (CCP).

## Author contributions

MH and EB conceived and designed the experiments. EB performed the experiments with assistance from SH for data collection and processing. MH and EB analysed the data and wrote the manuscript. All authors contributed to the writing and editing of the manuscript.

## Acknowledgements

EB gratefully acknowledges AstraZeneca for PhD funding. We thank Dimitri Chirgadze and Lee Cooper at the BiocEM Cryo-EM Facility (Department of Biochemistry, University of Cambridge) for their support and for access to Talos Arctica for collection of the dataset containing hPCC. We thank the Cambridge Pharmaceutical Cryo-EM Consortium (Department of Materials Science & Metallurgy, University of Cambridge) for access to and assistance with data collection on Titan Krios.

---

**Figure S1.**
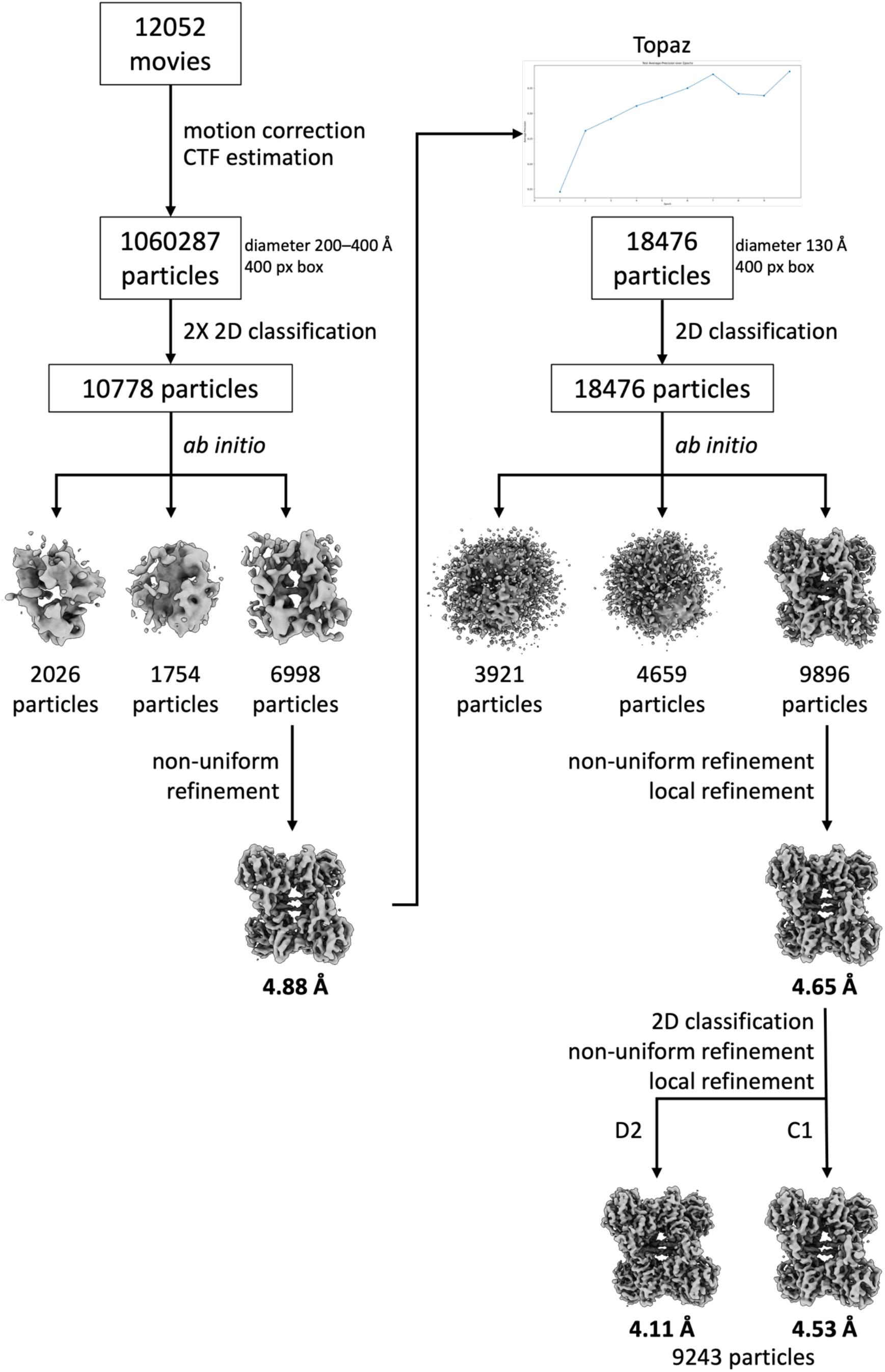
Cryo-EM data processing workflow for PRMT5:MEP50 complex. The data were processed in CryoSPARC. Volumes produced by *ab initio* reconstruction and subsequent non-uniform and local refinement are shown. Resolutions were evaluated at FSC = 0.143. After the initial volume was obtained from 6998 particles, a Topaz model was trained to search the dataset for further particles of the suspected contaminant, resulting in a final set of 9243 particles. The reported resolution of the volume was marginally improved when refined with D2 symmetry imposed.

